# Landscape and climatic features drive genetic differentiation processes in a South American coastal plant

**DOI:** 10.1101/2020.07.02.184366

**Authors:** Gustavo A. Silva-Arias, Lina Caballero-Villalobos, Giovanna C. Giudicelli, Loreta B. Freitas

**Affiliations:** Professorship for Population Genetics, Department of Life Science Systems, Technical University of Munich Freising, Germany; Laboratory of Molecular Evolution, Department of Genetics, Universidade Federal do Rio Grande do Sul, Porto Alegre, RS, Brazil

**Keywords:** *Calibrachoa heterophylla*, colonization, gene flow, genetic structure, landscape genetics, Solanaceae, South Atlantic Coastal Plain

## Abstract

**Background:** Historical and ecological processes shape patterns of genetic diversity in plant species. Colonization to new environments and geographical landscape features determine, amongst other factors, genetic diversity within- and differentiation between-populations. We analyse the genetic diversity and population structure of *Calibrachoa heterophylla* to infer the influence of abiotic landscape features on the level of gene flow in this coastal species of the South Atlantic Coastal Plain.

**Results:** The *C. heterophylla* populations located on early-deposited coastal plain regions show higher genetic diversity than those closer to the sea. The genetic differentiation follows a pattern of isolation-by-distance. Landscape features, such as water bodies and wind corridors, and geographical distances equally explain the observed genetic differentiation, whereas the precipitation seasonality exhibits a strong signal for isolation-by-environment in marginal populations. The estimated levels of gene flow suggest that marginal populations had restricted immigration rates enhancing differentiation.

**Conclusions:** Topographical features related to coastal plain deposition history influence population differentiation in *C. heterophylla*. Gene flow is mainly restricted to nearby populations and facilitated by wind fields, albeit without any apparent influence of large water bodies. Furthermore, differential rainfall regimes in marginal populations seem to promote genetic differentiation.

## BACKGROUND

Coastal areas in South America constitute distinct landscapes with unique abiotic and biotic compositions. Many geomorphological, climate, oceanographic features, and colonization events from the surrounding biomes shape the ecosystems of these areas [1–6]. Therefore, South American coastal flora shows a peculiar diversity of species, ecosystems, and various biogeographical processes shaping population demography [7, 8]. Although studies on plant diversification in South America have received increased attention, analyses focusing on the colonization of coastal areas, migration and gene flow between populations, and recent speciation events are still scarce [9–11]. There is indeed a lack of studies assessing the genetic diversity of wild plants from open areas such as sand dune and grassland plant communities from South-American coastal environments [12–14]. There are few works linking landscape genetics and phylogeography for coastal plant species in South America. Moreover, several gaps remain for fully interpreting population structure on spatially correlated genetic differentiation [15]. Disentangling the factors influencing gene flow is also important for understanding evolutionary dynamics at the extremes of species distribution [16] where processes such as local adaptation or peripatric speciation occur.

The species’ geographical distribution and genetic (nucleotide) diversity result from historical and contemporary processes acting together with ecological factors [17–19]. Colonization of new habitats and subsequent genetic isolation are critical events in the eco-evolutionary dynamics of coastal plant populations [20]. It is possible to reconstruct such events because the spread to new environments generates footprints on the genetic diversity and genetic spatial structure of populations [21]. Coastal regions have common environmental characteristics, such as intrinsic linear distributions, high salinity, wind strength, and tidal influence, which make these regions exciting models for studying genetic differentiation in response to climatic changes, changes in physical barriers, and in ecological features [22–24].

*Calibrachoa heterophylla* is a perennial nightshade shrub growing in dunes and sandy grasslands predominantly along the South Atlantic Coastal Plain (SACP). Previously phylogeographical assessment based on plastid markers supports that the species likely originated and subsequently diversified into four intraspecific lineages between 1 - 0.85 Mya [14]. These lineages have remained isolated by riverine barriers until their recent expansions following the formation of the SACP (400 – 7 kya), which determines their current geographical range. Currently, the species shows a continuous distribution along the SACP with a strong spatial genetic structure on the plastid markers albeit without conspicuous geographical barriers separating the intraspecific lineages [14]. This raises the questions whether nuclear polymorphism resembles the observed patterns in the chloroplast and if contemporary landscape features affect and shape the genetic structure of the species. We addressed these questions analysing a new set of polymorphic nuclear microsatellite markers and a comprehensive set of topographical and environmental variables in a spatial explicit framework.

The SACP is a flat, continuous, and open region constituting the most extensive coastal region in South America. The region extends NE-SW for approximately 600 km, is occupied mostly by large coastal lakes, and is crossed by two perennial water channels [25, 26]. This coastal formation gradually arose during sea-level transgressions and regressions caused by glacial-interglacial cycles during the last 400 ky. The most substantial transgression and regression cycles led to the formation of four main sand barriers that are positioned parallel to the coastline (barrier-lagoon systems I to IV; [25, 27]). Harsh environmental features such as strong spring-summer sea breezes from the Northeast and high insolation (solar irradiance) strongly influence the SACP [28] and consequently define suitable habitats for endemic plant species (e.g., [13, 14, 29]).

Currently, there are few studies assessing the genetic diversity of wild plants from the South American coastal environments (see [30, 31] for few examples). Even less works explicitly evaluate the relative influence of physical landscape (i.e., distance and topographical features) and environment on the genetic differentiation [13, 32]. Therefore, a landscape genetics assessment for *C. heterophylla* is useful to bring new insights to understand the evolution of coastal plants whereascomplementing the historical divergence processes described in *Petunia integrifolia* and *C. heterophylla* [13, 14, 29]. We hypothesize that the SACP colonization process altered the historical pattern of genetic structure shaped before the SACP deposition because the lineages came into contact due to the absence of strong geographic barriers along the SACP. Moreover, we expect that the geographical distance and differential features of the physical environment along the SACP, such as the age of the barrier-lagoon deposition, the presence of big water bodies, wind strength, and climatic gradients shaped local patterns of population admixture.

This study aims to understand the forces responsible for the current patterns of genetic structure in the coastal nightshade *C. heterophylla*. Based on an evaluation of relevant environmental and topographical features of the SACP and the analysis of polymorphic microsatellite markers, we (I) identify and infer the parameters of contemporary and historical factors promoting genetic divergence (colonization process, demographic process, rates of gene flow) and (II) assess which topographical and climatic factors determined the population differentiation and gene flow during the recent colonization of the SACP. We discuss the results in the light of relevant drivers of genetic diversification already identified for SACP species to find general scenarios shaping evolutionary trajectories of coastal plants in South America.

## RESULTS

### Genetic diversity

We found 140 alleles across ten microsatellite loci. The mean number of alleles per locus was 14, ranging from seven (Che59) to 17 (Che81). All loci showed higher *H*_e_ than *H*_o_ (Fig. S1) with 25% of the locus-population combinations showing a departure of HWE (P < 0.05). We detected a significant linkage disequilibrium signal (P < 0.01) for several loci pairs, however as the linkage pattern was not consistent across populations for any loci pair, we assumed linkage equilibrium and maintained all loci in the analyses. micro-checker analysis did not show evidence of null alleles, scoring errors, or stutter peaks for any locus.

Populations located outside of SACP (I1-3) and those collected around the Patos Lagoon (W1-2 and S1-2) showed higher genetic diversity (Fig. 1B). In contrast, the coastal populations located at the northern and southern edges of species distribution in SACP (N1 and S6) showed lower genetic diversity values. Average *H*_o_ values across loci ranged from 0.72 (I1) to 0.31 (N1) and for *H*_e_ from 0.74 (I3) to 0.48 (S3). We found 22% of the alleles restricted to a single population, with W1 showing the highest number of private alleles (eight), whereas W3, S1, S3, and S4 populations had no private alleles. Garza-Williamson values ranged from 0.39 (I2) to 0.83 (N2). We found positive and significant *F*_IS_ values for five populations (Table 1), all of them located at the borders (northern and southern) of species’ distribution inside the SACP (Fig. 1).

**Figure 1.**
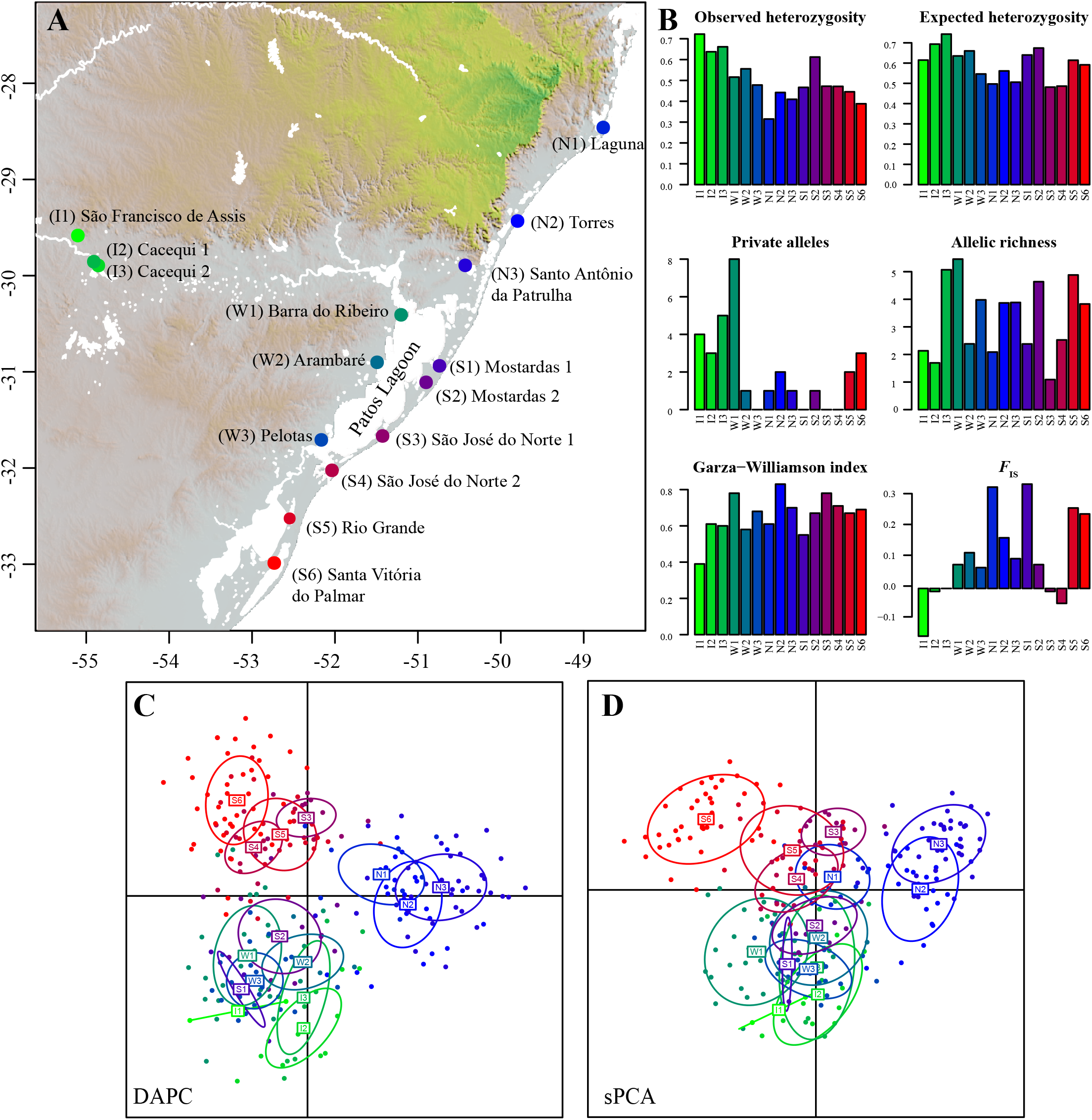
(A) Locations of *Calibrachoa heterophylla* populations. (B) Graphical representation of the mean genetic diversity statistics estimated for each population across all microsatellite loci. (C) Scatterplot of the DAPC analysis. (D) Scatterplot of the sPCA. Populations’ numbers and colors follow each panel.

**Table 1.** Sampling information and inbreeding coefficients for *Calibrachoa heterophylla* populations. Population ID codes follow Fig. 1A.

| Pop ID | N | Locality | Long | Lat | $F_{IS}$ |
| --- | --- | --- | --- | --- | --- |
| I1 | 4 | São Francisco de Assis | -55.10077 | -29.58307 | -0.16 |
| I2 | 10 | Cacequi 1 | -54.85375 | -29.8947 | -0.01 |
| I3 | 9 | Cacequi 2 | -54.90852 | -29.85478 | 0 |
| W1 | 27 | Barra do Ribeiro | -51.20255 | -30.40754 | 0.08 |
| W2 | 4 | Arambaré | -51.49195 | -30.90082 | 0.12 |
| W3 | 23 | Pelotas | -52.16478 | -31.70757 | 0.07 |
| N1 | 10 | Laguna | -48.76501 | -28.45991 | 0.34** |
| N2 | 23 | Torres | -49.79809 | -29.43227 | 0.17** |
| N3 | 37 | S Antônio da Patrulha | -50.42936 | -29.89291 | 0.10* |
| S1 | 3 | Mostardas 1 | -50.73934 | -30.93746 | 0.35 |
| S2 | 13 | Mostardas 2 | -50.90112 | -31.10909 | 0.08 |
| S3 | 12 | S José do Norte 1 | -51.42576 | -31.66673 | -0.01 |
| S4 | 11 | S José do Norte 2 | -52.03612 | -32.02393 | -0.05 |
| S5 | 26 | Rio Grande | -52.54661 | -32.52396 | 0.27*** |
| S6 | 41 | S Vitória do Palmar | -52.7323 | -32.98765 | 0.25*** |

\*P-value < 0.05; \*\* P-value < 0.01; \*\*\* P-value < 0.001

### Genetic structure

The recovered population structure showed a concordant geographic signal for marginal populations and higher admixed membership for populations located in geographical transitional regions (Fig. 2; Fig. S3). The best K = 4 was inferred from the ΔK method in Structure (Fig. S2A), whereas the DIC values from TESS showed the lowest standard for K = 2-4 runs and reach a plateau after maxK = 8 (Fig. S2B). DAPC showed the lowest BIC score for K = 8 (Fig. S4). Results obtained with all approaches showed consistent clustering of three to four well-differentiated groups.

**Figure 2.**
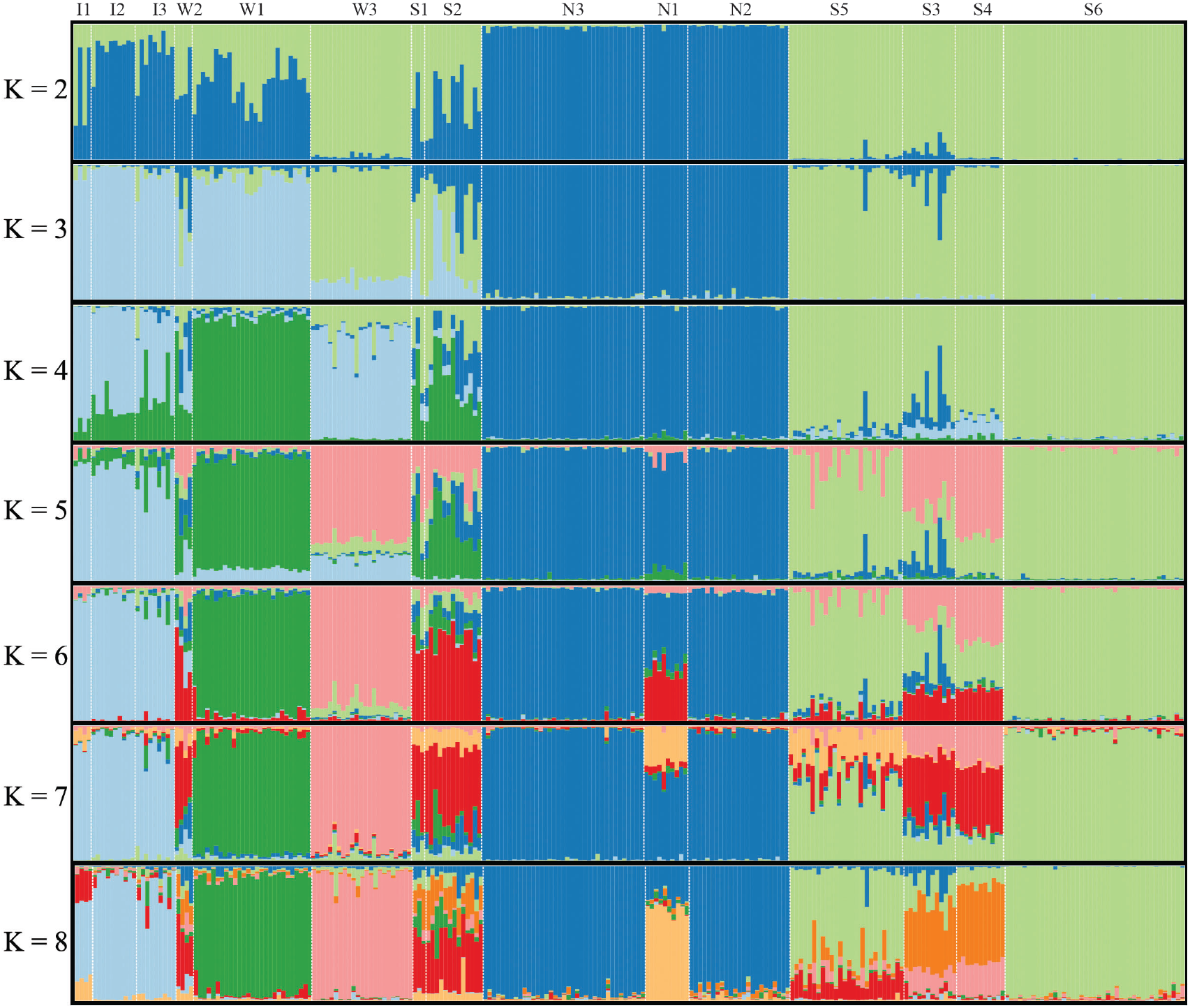
Bar plots of the individual membership for K = 2-8 genetic clusters as estimated with TESS. Populations are separated by white dashed lines and named on the top side.

Populations from the North of the SACP (N1-3) became the most differentiated group supported by the K = 2 clustering of Structure (Fig. 2) and TESS (Fig. S3) analyses, and the two main axes of both DAPC and sPCA (Fig. 1C,D). Considering three clusters, all approaches consistently recovered one group for the northern coastal populations (N1-3), a second group for the southern coastal populations (S3-6), and the third cluster for the three inland populations (I1-3) and the populations from the West side of the Patos Lagoon (W1-3). The two remaining populations (S1-2) showed a higher affinity with the Inland-West group in the exploratory analyses (Fig. 1C-D) and highly admixed memberships in the Bayesian clustering methods (Fig. 2, S3).

### Migration rates

The mean migration rate estimated with BayesAss was 0.015. However, only four population pairs showed higher posterior effective migration rates and confidence intervals above zero. Among them, the most outstanding was S2 -> W2 (*N*_m_ ≈ 0.08; 95% CI 0.01 - 0.14), supporting migration between populations separated by the Patos Lagoon. The other three cases involved neighbour populations, I2 -> I3 (*N*_m_ ≈ 0.12; 95% CI 0.05-0.19), S2 -> S1 (*N*_m_ ≈ 0.07; 95% CI 0.01-0.13), and S4 -> S3 (*N*_m_ ≈ 0.16; 95% CI 0.09-0.22). Migration estimates obtained from independent runs of BAYESASS showed similar values (Table S1).

The model-based coalescent approach implemented in migrate-n supported the *step-stone from coast* as the most likely historical migration model between population groups (Table 2; Fig. S5D). Parameter estimation showed that the ‘Inland’ group had the highest mean θ, which was around eight times higher than the θ estimated for ‘West’ and ‘North’ groups, and around 20 times higher than the θ estimated for the ‘South’ group (Table 2). Migration from ‘West’ to ‘Inland’ showed the highest mean M being two times higher than the ‘North’ to ‘West’ and five times higher than the ‘South’ to ‘West’ values (Table 2). We verified that all estimated parameter estimation procedures did reach convergence (effective sample > 10 000 and posterior estimates showed unimodal distribution; Additional file 1).

**Table 2.**
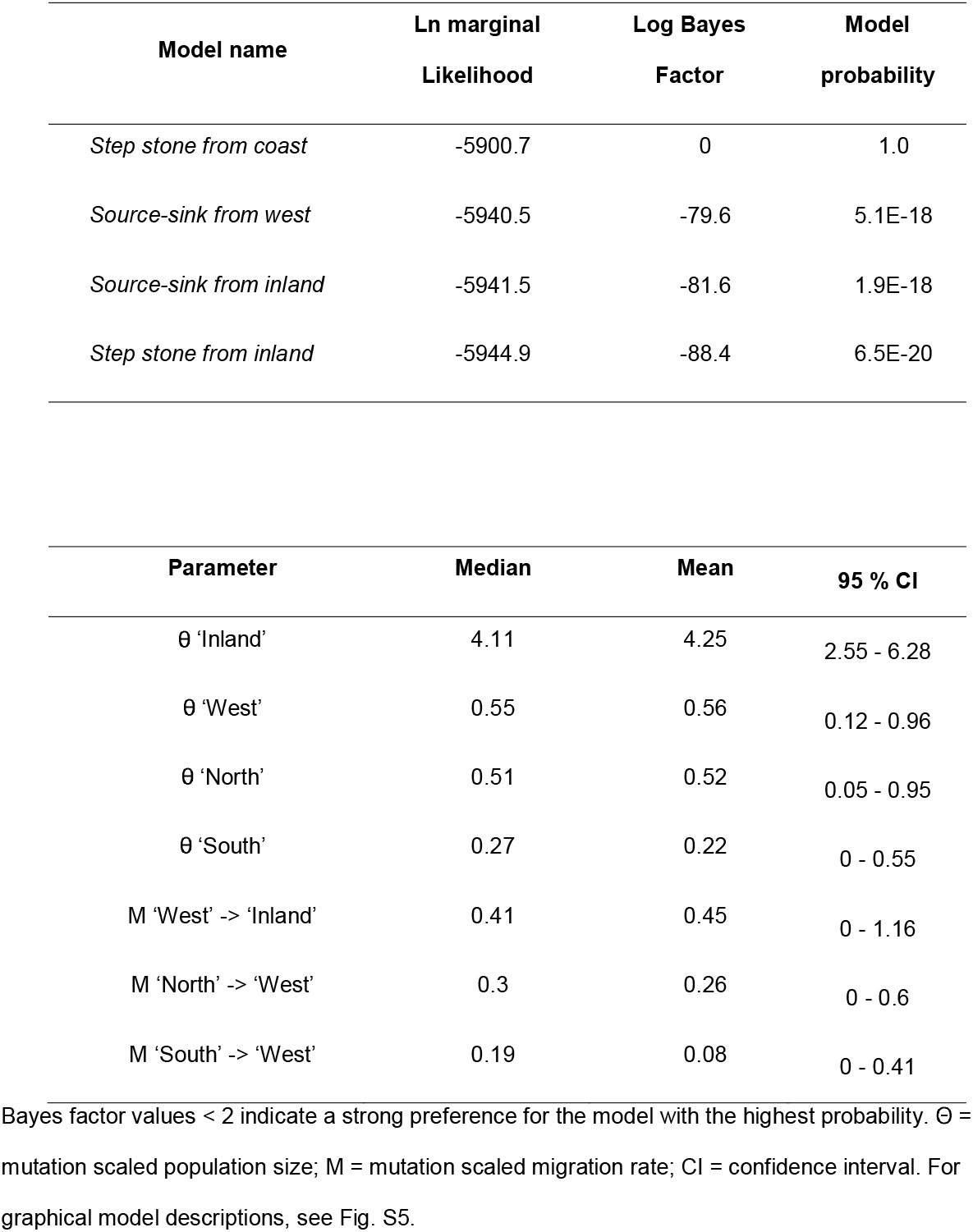
Model support statistics (upper panel) and parameter estimations taken from the best-supported model (lower panel).

### Isolation-by-distance, isolation-by-environment, and resistance tests

Measures of population differentiation *F*_ST_ ranged from 0.01 (S1 - S2 populations) to 0.54 (N1 - S3 populations; Fig. 3A). The Mantel test also supported a positive correlation between the genetic and geographical distance matrices (Mantel’s r = 0.38, P < 0.001). We then explored relevant landscape and climatic features throughout the SACP as potential determinants of genetic differentiation based on the MMRR approach.

**Figure 3.**
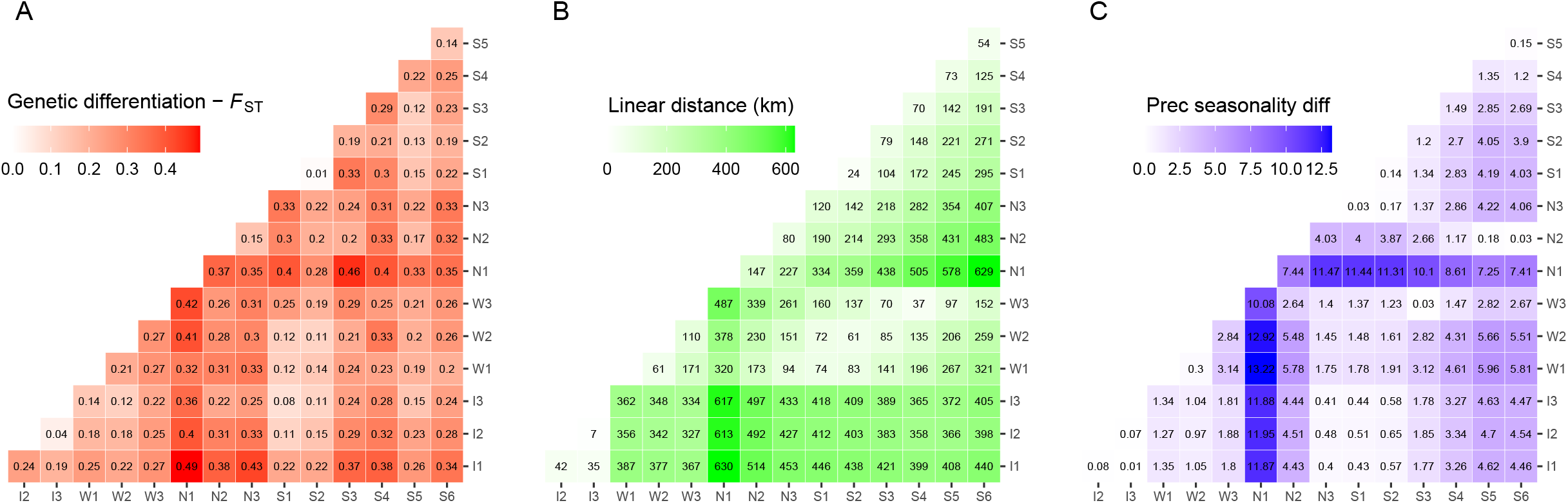
Interpopulation genetic, geographical, and environmental dissimilarity matrices. (A) Genetic differentiation index *F*_ST_; (B) Linear geographical distance; (C) Precipitation seasonality dissimilarity. A continuous color scale represents the values.

Further IBD tests assessing topographic cost distances models showed that the *continuous model* (landscape matrix with no topographic discontinuities) explained slightly better the genetic differentiation than the *water bodies model* (landscape matrix with water bodies as full barriers to population connectivity) (R^2^ = 0.16, β = 0.022, P = 0.006 and R^2^ = 0.14, β = 0.02, P = 0.011; respectively; Fig. S6). The relationship among climate variables and genetic differentiation including geographical distance showed significant association only for precipitation seasonality (Fig. 3C; R^2^ = 0.35, P = 0.003; β_precseason_ = 7.5 × 10-3, P = 0.05; β_Euc_ = 1.1 × 10-7, P = 0.02, respectively).

The “windscape” connectivity matrix accounting for both wind strength and direction measures (Figs. 4B, C) showed a North-to-South asymmetric step-stone pattern where marginal populations resulted strongly isolated and showing more intense wind influence at the central part of the SACP. Moreover, populations located at the West side of the Patos Lagoon became receptors from coastline populations. The coast distance wind matrix showed significant correlation with the *F*_ST_ genetic distance matrix (R^2^ = 0.19, β = 0.001, P = 0.0037; Fig. 4A).

**Figure 4.**
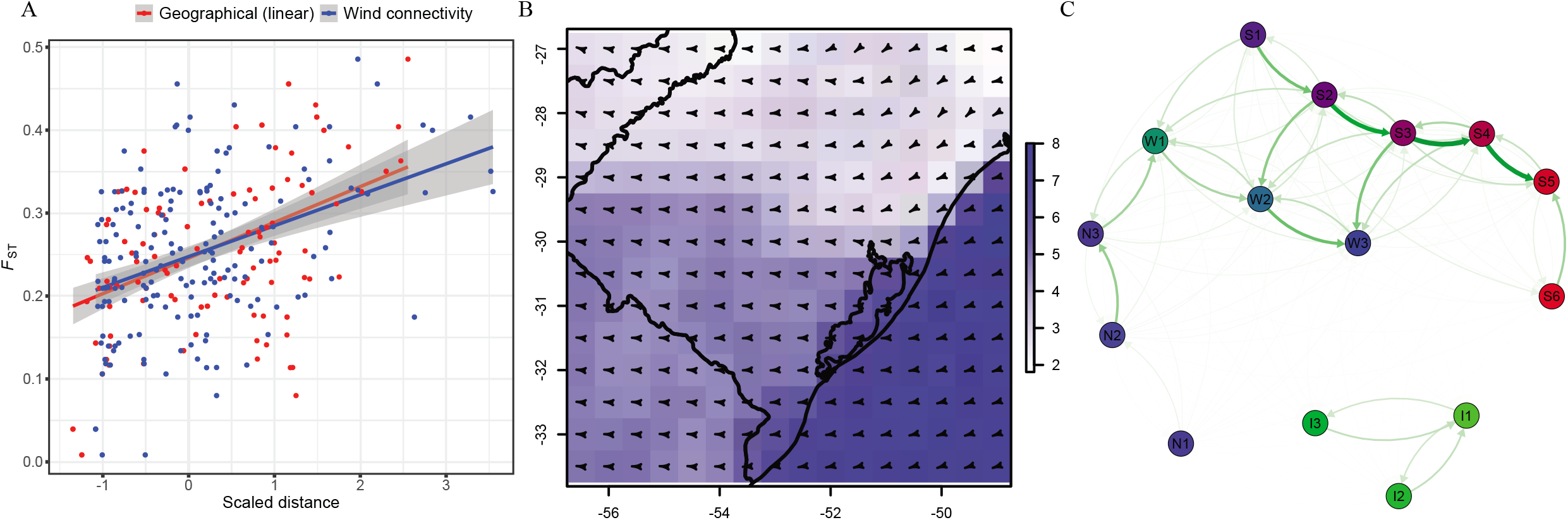
Spring season wind fields in the South Atlantic Coastal Plain and inference of wind in the population connectivity and gene flow. (A) Correlation between geographical distance (red) and wind connectivity (blue) cost distance matrices with the *F*_ST_ genetic distance matrix; (B) Mean values 2011-2016 of wind speed (blue scale m/s) and direction (arrows) for the spring season (September-November); (C) Inter-population wind connectivity network.

## DISCUSSION

In this study we analyse the genetic diversity and structure of *Calibrachoa heterophylla* to infer the influence of topographical and environmental features on the population differentiation during the recent colonization of a coastal region in South America. The results support both contemporary and historical factors promoting genetic divergence throughout populations of a coastal plant species. Here, we provide consistent evidence for limited and asymmetric gene flow, mainly restricted by geographical distance. The populations from northern and southern edges of the species distribution show negligible historical and contemporary immigration rates related to historical and geographical isolation. We also found that one of the most outstanding topographical feature in the SACP, namely the large water bodies, does not constrain *C. heterophylla* populations’ gene flow. Gene flow seems promoted by the wind, at least between adjacent populations from the central portion of the SACP. Our results highlight the importance of considering both the physical landscape (contemporary) and phylogeographical context (historical) processes for complete interpretation of genetic differentiation processes.

### Role of historical, spatial, and environmental factors on the genetic differentiation in *Calibrachoa heterophylla*

There is a hierarchical pattern of genetic structure related to both historical and contemporary landscape features. The main clustering pattern mirrors the phylogeographical structure of *C. heterophylla* previously recovered with plastid markers [14]. The retention of historical signals of genetic structure in highly variable markers, such as microsatellites, is expected for studies involving the entire geographic range of species, reinforcing the importance of considering the historical patterns for interpreting landscape genetic analyses [33]. Moreover, northernmost populations from the ‘South’ group (S1 and S2) or the southernmost or the ‘West’ group (W3), given their intermediary location, display higher admixture values supporting secondary gene flow between previously differentiated intraspecific lineages (Fig. 1).

The influence of geographical distance on the genetic structure is evident in the genetic structure of *C. heterophylla*. As expected, the effect of geographical isolation is stronger in peripheral populations such as S6, I1-3, and N1. Therefore, genetic drift due to long-term geographical isolation mainly explains the strong differentiation at the edges. The immigration of populations at the SACP edges falls within the lowest estimates (Fig. S3). However, differential conditions at the edge of the distribution could also be involved. For example, the northern portion of the SACP presents significant differences in precipitation seasonality because of the influence of orographic rainfalls during the spring and summer seasons. This environmental feature significantly correlates with high genetic differentiation in northern populations (Fig. 3). These results point to a genetic divergence process enhanced by local adaptation. Ecological differentiation can promote selection against immigrants (maladaptive gene flow), leading to reduced gene flow, reproductive isolation, and enhancing the stochastic effects of genetic drift [34–36]. As this pattern is also seen in co-distributed coastal populations of *Petunia integrifolia* [13], further research is worthwhile to uncover potential convergence local adaptation processes related to precipitation differences.

### Environmental and geomorphological processes around the Patos Lagoon led to a secondary contact between previously diverged lineages

Intricate spatial and environmental influences on the genetic structure is exemplified through the discordant clustering patterns of population W3 with ‘South’ and ‘West’ groups depending on the approach. This feature reflects the intermediary geographic location between those two regions but also the fact that this zone shows fluctuating inland and coast environmental conditions. Both inter-annual rainfall differences and long-term climatic fluctuations, such as El Niño phenomenon, affect the fluvial discharge and wind currents responsible for the salinization and desalination processes in the Patos Lagoon [37, 38]. This environmental dynamic could periodically change the individuals’ establishment or survival rates of either coastal or continental gene pools probably leading to a mixed genetic pool in this region.

The populations W2-3 and S1-5, all located around the Patos Lagoon (Fig. 1A), show high levels of genetic admixture (Fig. 2; Fig. S1) and the lowest *F*_ST_ values (Fig. 3A). These results are consistent with the recent geomorphological history of the SAPC. During most of the Quaternary Period, two rivers (Jacuí and Camaquã), including several channels corresponding to their dynamic delta systems, maintained distinct inlets on the Patos Lagoon area [14, 39]. Only after the formation of the barrier systems III and IV (the most recent and closer to the shoreline coastal strips) between 12-7 kya, the Patos Lagoon reached its current conformation, and the current continuous coastline was established [40]. In contrast, the northern and southern regions, corresponding to the older barrier systems I and II (cf. Fig. 1 in [27], let to earlier expansion and differentiation of the coastal lineages that, later, spread and experienced a secondary contact on the East side of the Patos Lagoon generating the current patterns of genetic admixture in this region. The recent admixture processes are also supported by the lack of private alleles in W3, S1, S3, and S4 populations (Fig. 1B). This geomorphological history seems to determine common patterns among co-distributed species from the region. Despite the differences in divergence times of the intraspecific lineages of *C. heterophylla* (earlier) and the coastal populations of *P. integrifolia* (recent) [14, 29], these co-distributed taxa share the patterns of high genetic admixture in populations located at the East side of Patos Lagoon [13].

The East side of Patos Lagoon (seashore side) undergoes the strongest wind influence within the SACP (Fig. 4B; [41]), potentially increasing secondary seed dispersal alongside the region generating, in consequence, higher admixture rates. The gene flow estimations among *C. heterophylla* populations support an asymmetric migration from coastal to inland locations, even at long distances crossing the coastal lakes (Figs. 1 and 4). Although wind can significantly influence coastal environments, and it shapes large-scale population differentiation, gene flow, and genetic diversity [42], the influence of wind variables is poorly explored in landscape genetics approaches. Wind conditions also affect population dynamics of Tuco-Tuco rodents (*Ctenomys* sp.) in the SACP [43]. This convergent factor between co-distributed taxa supports that the current dynamics in the topographical and environmental conditions in SACP play a role in the structuration at the community level.

Our findings expand the knowledge of genetic differentiation and diversification processes across coastal areas. According to Wieringa et al. [44], our study highlights multiple processes likely influencing genetic structure. For example, *C. heterophylla* and the coastal lineage of *P. integrifolia* have strong differences in the divergence times and intraspecific differentiation but convergent contemporary distribution and genetic structure. Our results suggest that a complex mixture of features related to physical barriers, geographic distance, and environment along the SACP shape shared contemporary genetic differentiation patterns on the region’s species. Therefore, considering historical and recent diversification processes is crucial to interpret either shared or idiosyncratic patterns in contemporary genetic structure. We strongly encourage new research into the environmental factors driving genetic structure within and among populations on plant species distributed along different coastal regions from South America.

## CONCLUSIONS

*Calibrachoa heterophylla* recently colonized the SACP leading to a typical linear distribution shape for coastal species. The species shows limited and asymmetric gene flow patterns, mainly influenced by geographical distance and wind. The presence of big water bodies, which constitutes the most outstanding topographical feature in the SACP, does not constrain inter-population gene flow. Negligible historical and contemporary immigration rates in marginal populations coupled to contrasting precipitation conditions could promote genetic differentiation in the northern and south marginal populations. Recent admixture from previously differentiated populations and higher gene flow explain the genetic diversity in the most recently formed coastal areas and more substantial wind influence region of the SACP. Our results highlight the need to integrate both phylogeographic and landscape genetic approaches to disentangle processes affecting the genetic differentiation of coastal plant species.

## METHODS

### Study system

The species of *Calibrachoa* (Solanaceae) occur in subtropical and temperate grasslands in southern Brazil, northeast Argentina, and Uruguay. The genus encompasses ca. 30 species, among which *C. heterophylla* is the only species that colonized coastal environments [45]. This species is diploid (2n = 18), semi-prostrated, and displays purplish bee-pollinated flowers; the fruits are capsules and produce dozens of tiny seeds (< 1.4 mm) with no dispersal mechanisms. *C. heterophylla* occupies open sandy grasslands, dunes, or rocky outcrops in lakeside or marine environments from ~ 28 Lat S to 32 Lat S in the SACP [14]. Longitudinally, populations of *C. heterophylla* occur from the seashore to less than 90 km from the coast, with few populations separated from the seashore by big lagoons. Just one disjointed and small population group occurs outside SACP, restricted to the sandbanks alongside the Santa Maria River basin, ~ 55 Long W (Fig. 1A).

### Sample collections

For this study, we used all samples included in Mäder et al. [14] plus additional samples from the wild for a total sampling of 253 individuals from 15 locations (hereafter populations; Fig. 1A) that covered the entire species’ distribution. We collected leaves of all individuals found in each locality and preserved them in silica gel. The number of individuals per population varied from three to 41 (Table 1). We also sampled complete branches for herbarium specimens from those localities subsequently deposited in BHCB (Universidade Federal de Minas Gerais, Belo Horizonte, MG, Brazil) and ICN (Universidade Federal do Rio Grande do Sul, RS, Brazil). Plant identification was performed by G. Mäder, J. Fregonezi, or G. Silva-Arias and then confirmed by the group expert J. R. Stehmann. We required no specific permits since collection localities correspond to neither private properties nor protected areas. Also, the field studies did not involve endangered or protected species. This work was conducted under MP 2.186-16 of the Brazilian Federal Government.

### Laboratory procedures and genotyping

The total DNA was extracted following a CTAB-based protocol [46] and amplified for ten anonymous microsatellite loci developed for *C. heterophylla* (Che18, Che59, Che119, Che26, Che34, Che81, Che82, Che85, Che72, and Che126) following optimized protocols for PCR and genotyping procedures [47]. We used micro-checker [48] to estimate genotyping errors due to stutter bands, allele dropout, or null alleles.

### Characterization of the genetic diversity

We performed tests for linkage disequilibrium and deviations from HWE within each population for each locus. We assessed the significance of HWE deviations using 10^6^ Markov chain steps and Fisher’s exact probability tests in Arlequin v.3.5 [49]. We estimated the genetic diversity based on average rarefied allelic richness, private alleles, *H*_o_, *H*_e_, the G-W index, and *F*_IS_ (with confidence limits from 1000 bootstrap resampling over loci) using the poppr v.2.8.5 [50] and hierfstat v.0.04-22 [51] packages in R v.3.6.3 package [52], and Arlequin.

### Population genetic structure

We assessed the genetic structure employing two model-based clustering methods and two exploratory data analyses [53]. The model-based clustering methods used are Structure v.2.3.4 [54] and the spatial Bayesian clustering program TESS v.2.3 [55, 56]. These analyses provide estimates for the K ancestral clusters assuming HWE equilibrium, individual assignment probabilities and compute the proportion of each individual’s genome assigned to the inferred clusters.

For Structure analysis, the number of clusters evaluated ranged from 1 to the total number of populations (15), with ten independent runs per K-value. We performed each run using 2.5 × 10^5^ burn-in periods and 1.0 × 10^6^ Markov chain Monte Carlo repetitions after the burn-in, under an admixture model, assuming correlated allele frequencies [57], including *a priori* sampling locations as prior (*locprior*) to detect weak population structure. The *locprior* option is not biased toward detecting structure when it is not present and can improve the Structure results when implemented with few loci [58]. To obtain the K value that better explains the structure based on the genetic dataset, we assessed the measures of the ΔK method [59] that is useful to recover the hierarchical highest level of genetic structure.

TESS implements a spatial assignment approach to group individuals into clusters accounting for samples’ geographical locations, giving them higher probabilities of belonging to the same genetic cluster to those that are spatially closer in the connection network. For TESS, we ran100 000 generations, with 50 000 generations as the burn-in, using the CAR admixture model, and starting from a neighbor-joining tree. We ran 20 iterations for each value of maxK ranging from 2 to 15. We added a small perturbation to the original population coordinates with a standard deviation equal to 0.2 to obtain single different coordinates for each individual. We assessed the convergence inspecting the post-run log-likelihood plots and obtained the support for alternative K values inspecting the statistical measure of the model prediction capability from DIC [60]. We computed and plotted the average of DIC values to detect maxK value at the beginning of a plateau. Replicated runs of best K results for Structure and TESS were summarized and plotted with the Pophelper [61] R package.

Additionally, to detect genetic structure, we implemented the exploratory multivariate methods Discriminant analysis of principal components (DAPC [62]) and the spatial Principal Components Analysis (sPCA [63]) implemented in the Adegenet v.2.1.3 [64] R package. For the DAPC analysis, the SSR data were first transformed using PCA and keeping all PCs. The number of clusters that maximizes the between-group variability using the BIC score was optimized using the function *find.clusters*. To avoid overfitting, we set an optimal reduced number of PCs using the function *optim.a.score*.

The sPCA incorporates spatial information to maximize the product of spatial autocorrelation (Moran’s I) and the variance for each eigenvector, producing orthogonal axes that describe spatial patterns of genetic variation. The spatial information is included in the analysis using a spatial weighting matrix derived from a connection network. To test the effect of the neighbors definition on the results, we ran the sPCA using six different connection networks available in the function *chooseCN*. For this analysis, we used the same perturbed coordinates used in TESS analysis. Monte Carlo simulations (global and local tests) were used with 10 000 permutations to test for non-random spatial association of population allele frequencies for all implemented sPCA. Clustering patterns recovered with the DAPC and sPCA were visualized in scatter plots obtained with the function *s.class* in R.

### Historical and contemporary gene flow estimations

Contemporary asymmetric migration rates were estimated using the Bayesian approach implemented in BayesAss v.3.0 [65]. We ran 10^8^ iterations and a burn-in of 10^7^. We adjusted the mixing of allele frequencies, inbreeding coefficients, and migration rates parameters to 0.6, 0.6, and 0.3, respectively, to obtain acceptance rates of around 40%. We assessed convergence by examining the log-probability plots and the effective sample sizes for each run using Tracer v.1.6 [66] and looking for consistency of the migration estimates among three independent runs with different initial seed numbers.

We assessed historical gene flow by testing the support of four alternative scenarios given our genetic dataset using Bayes factors calculated from the Bézier log-marginal likelihood approximations [67]. We used the coalescent-based Migrate-N v.3.2.6 [68] software to estimate the mutation-scaled effective population size (θ) and the mutation scaled migration rate (M) parameters. We pooled the populations into four groups for all models according to the geographical distribution and genetic structure (see Results Figs. 1A and 2). The ‘Inland’ group included the I1-3 populations; the ‘West’ group encompassed the W1-3; ‘North’ included the N1-3 populations, and the ‘South’ group clustered the S1-6 populations.

We evaluated four migration models: (1) *source-sink from inland* with unidirectional migration from ‘Inland’ group to the remaining groups; (2) *source-sink from the West* with unidirectional migration from ‘West’ group to the remaining groups; (3) *step-stone from inland* with unidirectional migration from Inland to West and from West to North and South; and (4) *step-stone from coast* with unidirectional migration from North to West, from South to West, and from West to Inland (Fig. S5).

We ran the Migrate-N Bayesian inference in the Cipres Science Gateway v.3.3 [69], with one long chain of 5 × 10^6^ steps, sampling at every 100^th^ increment, and a burn-in of 3 × 10^4^ steps. We used uniform priors and slice sampling for both θ and M ranging from 0 to 20 (mean = 10, delta = 0.5). We used a heating scheme MCMCMC with four parallel chains and temperatures of 1, 1.5, 3, and 10^6^.

### Space, topography, environment, and genetic differentiation

Spatial correlation patterns under IBD generate bias in several genetic structure tests [15, 70, 71]. Therefore, we assessed the IBD through linear regression of linearized pairwise *F*_ST_ genetic distances and log-transformed geographical distances [72] using a Mantel test, assessing the significance with 10 000 randomizations in Vegan v.2.5-6 [73] R package. Pairwise *F*_ST_ [74] matrix was calculated with the Hierfstat package and geographical inter-population distance matrix by calculating the linear Euclidean distance between X and Y UTM 22S (reference EPSG: 32722) populations’ coordinates transformed from Long/Lat coordinates with Rgdal v.1.0-4 [75] R package.

We tested IBE models to examine whether differences in climatic conditions explain inter-population genetic differentiation in *C. heterophylla*. Pairwise climatic dissimilarity matrices were obtained for the following bioclimatic variables: total annual precipitation, total annual days with rain, precipitation seasonality, mean annual temperature, mean summer maximum temperature, mean winter minimum temperature, mean temperature range, and temperature seasonality. Climatic data derive from raster layers specifically developed for the SACP obtained from a high-density sampling of climate stations throughout the region, geostatistical modeling, and spatial interpolation, as described in Silva-Arias et al. [13].

We also included a wind connectivity matrix in the IBE tests to evaluate the influence of strong winds in the SACP on the population migration rate. We calculated surface wind direction and speed data for the Southern Hemisphere’s spring months (September to November) 2011–2016 sampled every three hours. We downloaded the data from the Global Forecasting System using the rWind v.1.1.5 [76] R package. We transformed direction and speed values into raster layers for each sampled time using the *wind2raster* function to obtain transition layers using the function *flow.dispersion*. Finally, we calculated pairwise cost distance matrices with the function *costDistance* in gdistance v.1.3-1 [77] R package. We then averaged the matrices for the all-time series. We plotted the final matrix with the qgraph v.1.6.5 [78] R package.

We extended the IBD analyses using raster grids to test for possible models of inter-population differentiation linked to landscape discontinuities alongside the SACP. We outlined two coast distance models (Fig. S6): (1) the *continuous* (or null) model wherein no landscape discontinuity affects the interpopulation connectivity. We created a raster grid with all cells values equal to 1, including all cells on freshwater surfaces. This model is expected to resemble a Euclidean geographical distance, but it is more appropriate for comparisons with models based on circuit theory; and (2) the *water bodies* model, representing the widespread freshwater bodies in the SACP as connectivity barriers between populations. For that, we created a raster grid with all land cells values equal to 1, and cells within freshwater surfaces as complete barriers (*no data*). We generated pairwise cost distance matrices using the function *transition* in GDISTANCE package considering an eight-neighbors cell connection scheme, Long/Lat coordinates per population as nodes, and raster resolution of 0.09 degrees (~ 10 km).

We examined the relationships between *F*_ST_ and geographical or topographical distances (IBD) and environmental dissimilarity (IBE) using MMRR; [79] implemented in R.

## Supplementary Information

Fig. S1: Comparison observed *vs*. expected heterozygosity per microsatellite locus.

Fig. S2: Evaluation of the best number of clusters for Structure and TESS analyses.

Fig. S3: Bar plots of the individual membership obtained with Structure.

Fig. S4: Bayesian information criterion assessed with Discriminant Analyses of Principal Components.

Fig. S5: Graphical representation of the coalescent migration models.

Fig. S6: Graphical representation of the raster layers for topographic tests.

Table S1: Migration estimates obtained with three independent runs of BayesAss. Additional file 1. Migrate-n detailed output for the best-supported model showing parameter estimation values and convergence statistics.

## Abbreviations

BIC: Bayesian Information Criterion
CAR: conditional autoregressive
Che: *Calibrachoa heterophylla*
CI: confidence interval
cpDNA: plastid DNA
CTAB: cetyl-tetramethyl ammonium bromide
DAPC: Discriminant Analysis of Principal Components
DIC: deviance information criterion
*F*_IS_: inbreeding coefficient
*F*_ST_: genetic differentiation index
G-W: Garza-Williamson index
*H*_e_: expected heterozygosity
*H*_o_: observed heterozygosity
HWE: Hardy-Weinberg equilibrium
IBD: isolation-by-distance
IBE: isolation-by-environment
K: number of genetic clusters
km: kilometre
ky: thousand years
kya: thousand years ago
Lat: latitude
Long: longitude
M: mutation-scaled migration parameter
MCMCMC: Metropolis-coupled Markov Chain Monte Carlo
MMRR: multiple matrix regressions with randomization
NE: northeast
*N*_m_: Effective migration rate
PCs: principal components
PCR: polymerase chain reaction
R^2^: coefficient of determination’ r - correlation coefficient
SACP: South Atlantic coastal Plain
sPCA: spatial Principal Component Analysis
sPC: spatial Principal Components
S: south
SW: southwest
SSR: microsatellite locus
subsp.: subspecies
UTM: Universal Transverse Mercator coordinate system
W: west
θ: mutation-scaled population size

## Declarations

### Ethics approval and consent to participate

Not applicable.

### Consent for publication

All authors reviewed the manuscript and agreed with this publication.

### Availability of data and materials

All data generated or analyzed during this study are included in this published article [and its supplementary information files].

### Authors’ contributions

G.A.S.-A. and L.B.F. designed the study; G.A.S.-A. performed laboratory experiments, analyzed data, and interpreted the results; G.A.S.-A. and L.C.-V. discussed the results and wrote the manuscript; G.C.G. supported laboratory genotyping; L.B.F. supervised the project and provided resources for the research.

## Acknowledgements

The authors acknowledge G. Mäder and J. Fregonezi for help in fieldwork, J.R. Stehmann for plant identification, and the Technical University of Munich Publishing Fund for covering the costs for the publication of this article.

## SUPPLEMENTARY MATERIAL LEGENDS

**Figure S1**. The plot of observed *vs*. expected heterozygosity for each locus.

**Figure S2**. (A) Plots of the best K estimates for Structure results. (B) Plots of the mean and standard deviation of the deviance information criterion (DIC) obtained for each maxK assessed with TESS.

**Figure S3.** Bar plots of the individual membership for each genetic cluster obtained with Structure. White dashed lines separate populations, and names are indicated on the figure top side.

**Figure S4.** The plot of Bayesian information criterion (BIC) values obtained for each K number assessed using the multivariate method Discriminant Analyses of Principal Components.

**Figure S5.** Graphical representation of the four coalescent migration models tested in Migrate-N for *Calibrachoa heterophylla*. (A) Source-sink from inland; (B) Source-sink from the west; (C) Step-stone from inland; (D) Step-stone from coast.

**Figure S6**. Graphical representation of the raster layers used to calculate the connectivity values in topographic tests (A) Continuous model; (B) Water bodies model.

**Table S1.** Migration estimates obtained with three independent runs of BAYESASS. The values indicate the estimated posterior mean effective migration rate per generation [the fraction of individuals in population *i* (rows) that are migrants derived from population *j* (columns)], and the numbers in parentheses show the standard deviation. Bold values indicate the diagonal (intra-population estimates), and red values indicate the highest migration estimates (those with above zero 95% confidence intervals).

**Additional file 1.** Migrate-n detailed output for the best-supported model showing parameter estimation values and convergence statistics.

