## Supplementary_material for "Landscape and climatic features drive genetic differentiation processes in a South American coastal plant"

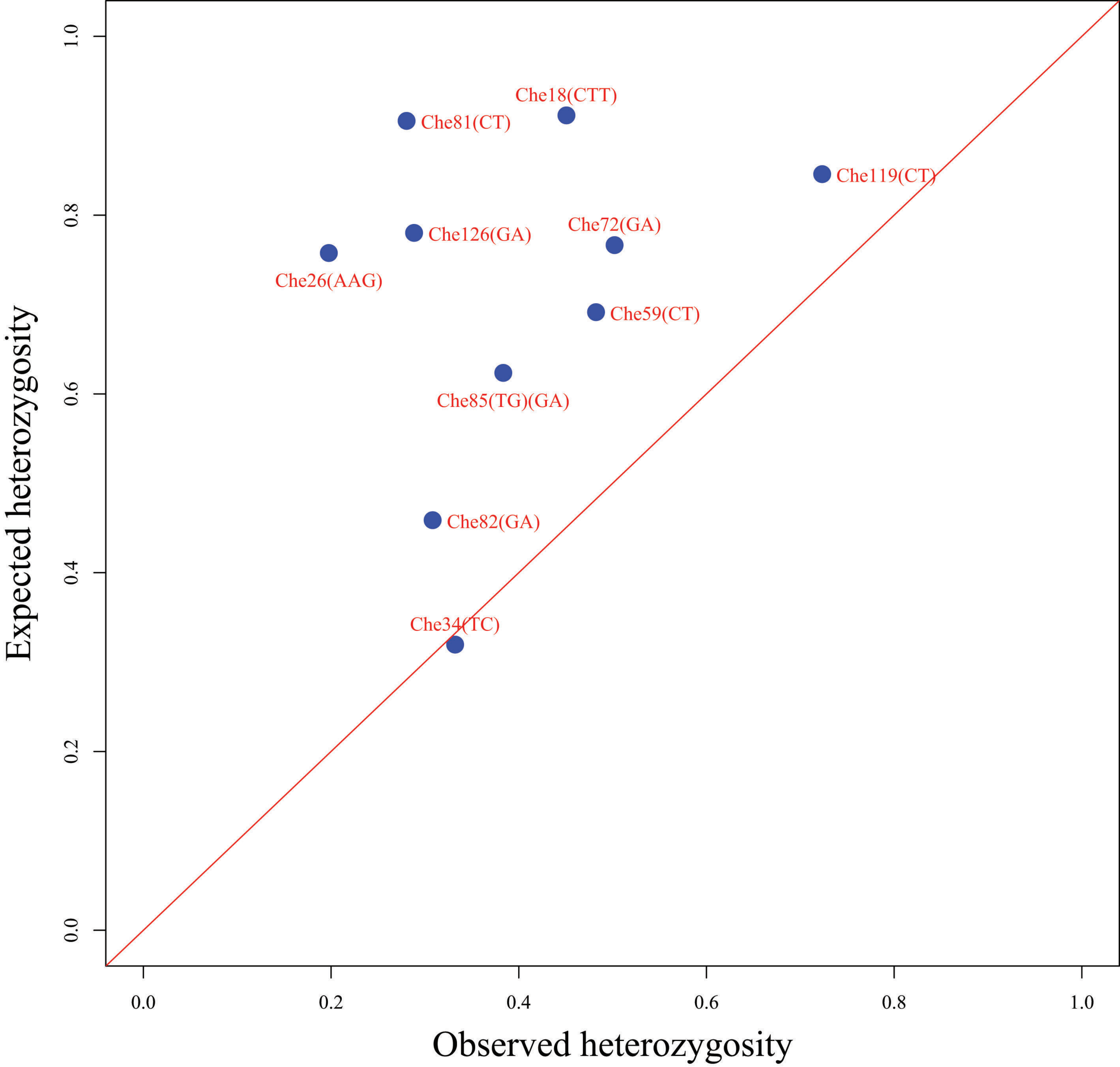

Che18(CTT)
Che81(CT)
Che119(CT)
Che126(GA)
Che72(GA)
Che26(AAG)
Che59(CT)
Che85(TG)(GA)
Che82(GA)
Che34(TC)
0.0
0.2
0.4
0.6
0.8
1.0
Expected heterozygosity
Observed heterozygosity

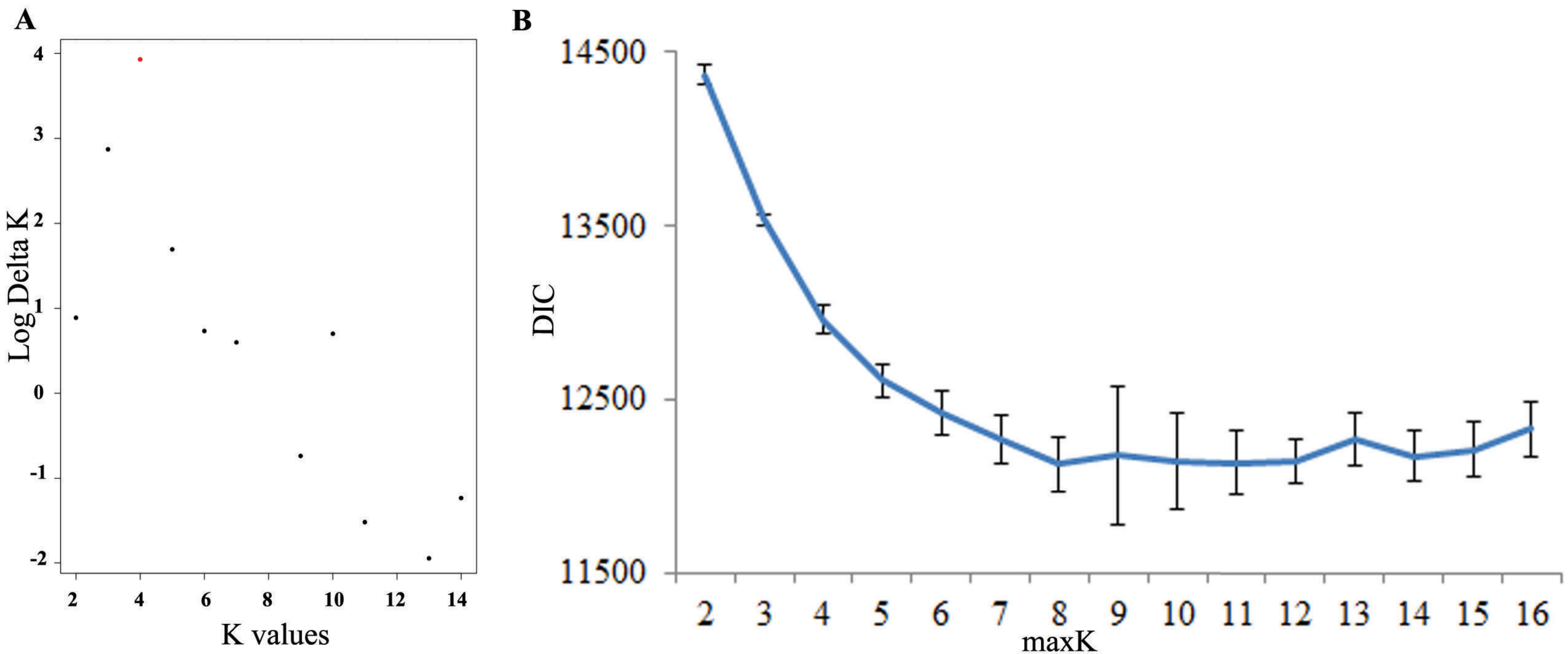
A
Log Delta K
K values
4
3
2
1
0
-1
-2
2
4
6
8
10
12
14
B
DIC
maxK
14500
13500
12500
11500

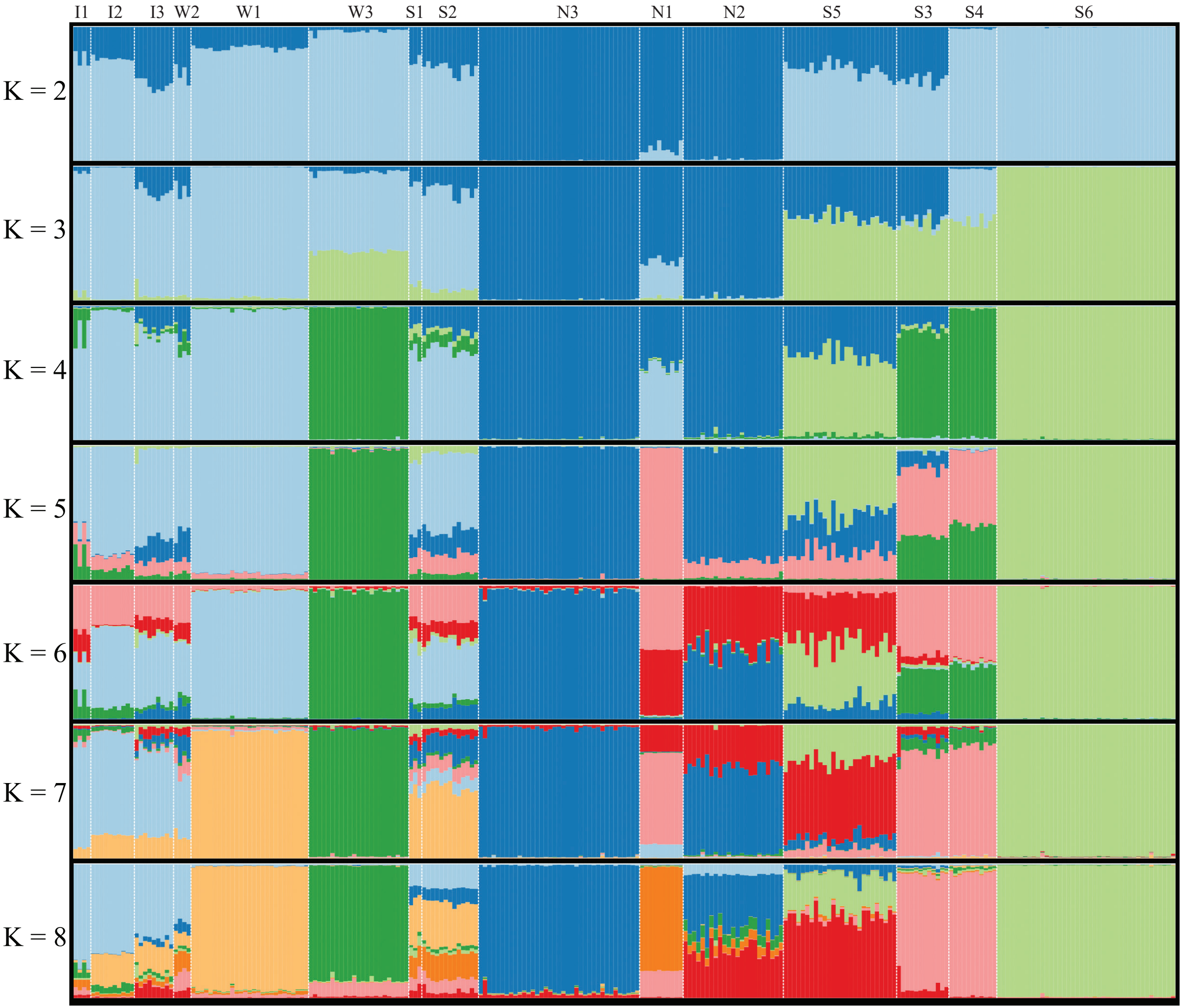
I1
I2
I3
W2
W1
W3
S1
S2
N3
N1
N2
S5
S3
S4
S6
K = 2
K = 3
K = 4
K = 5
K = 6
K = 7
K = 8

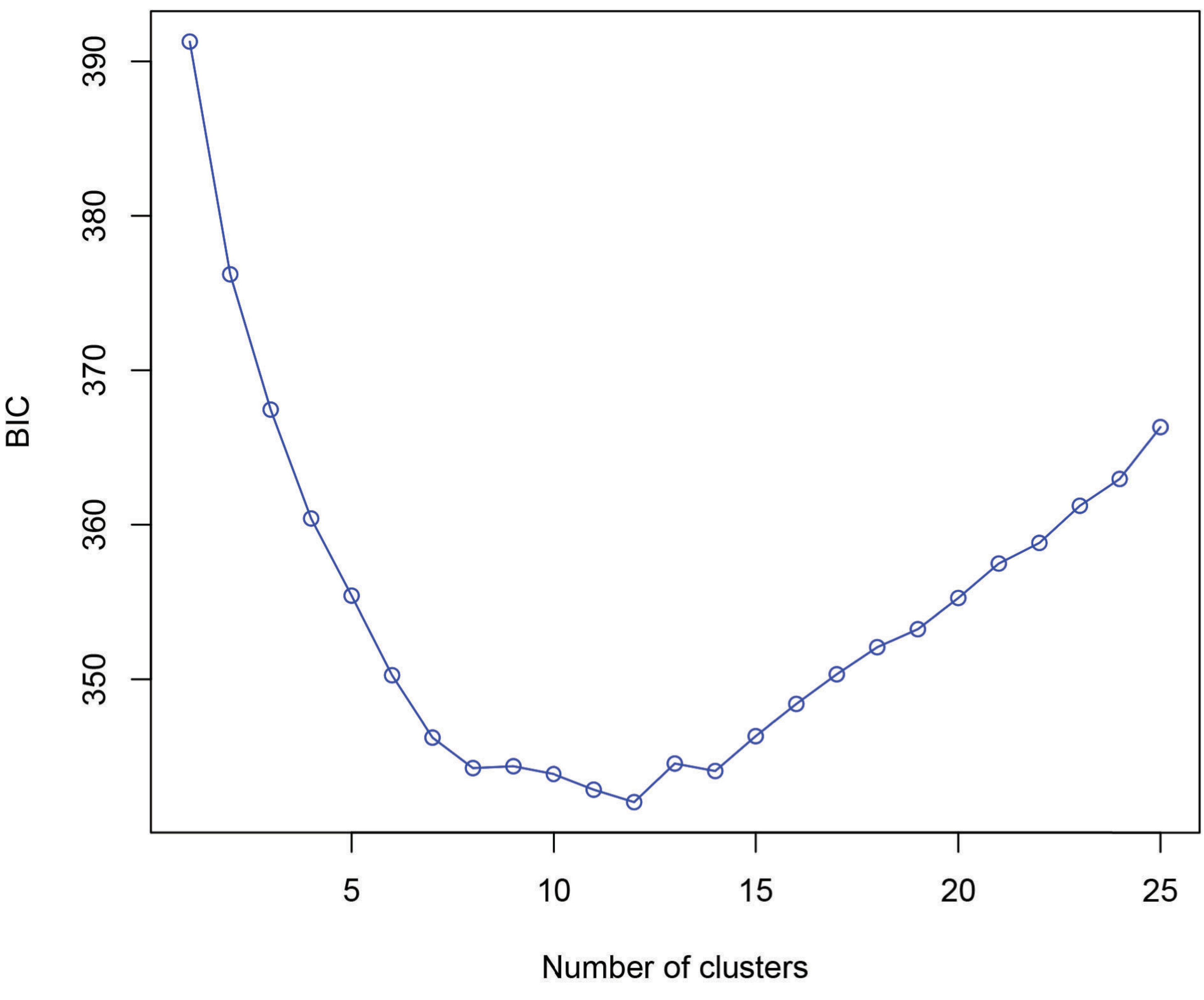

390
380
370
360
350
BIC
5
10
15
20
25
Number of clusters

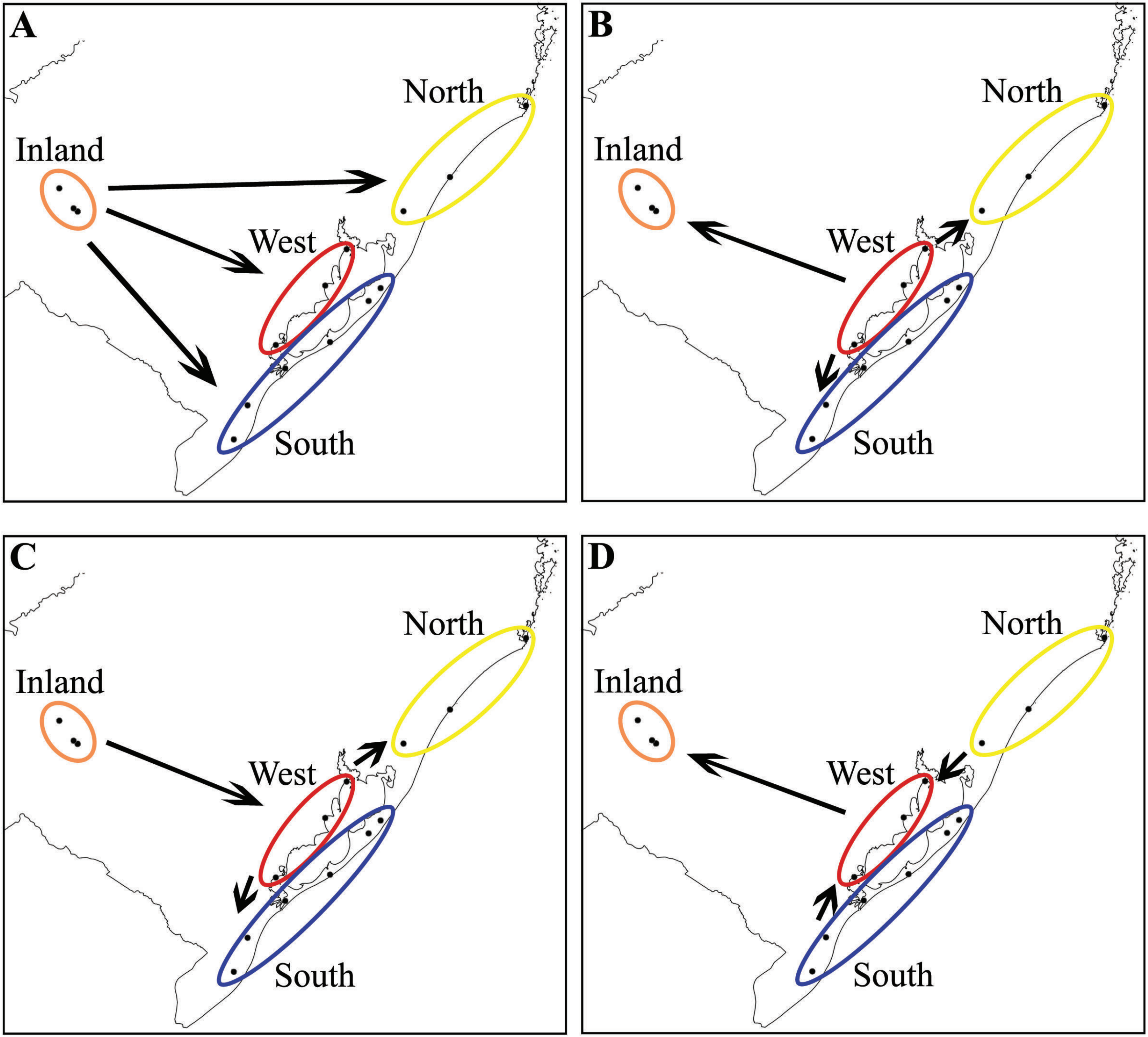
A
North
Inland
West
South
B
North
Inland
West
South
C
North
Inland
West
South
D
North
Inland
West
South

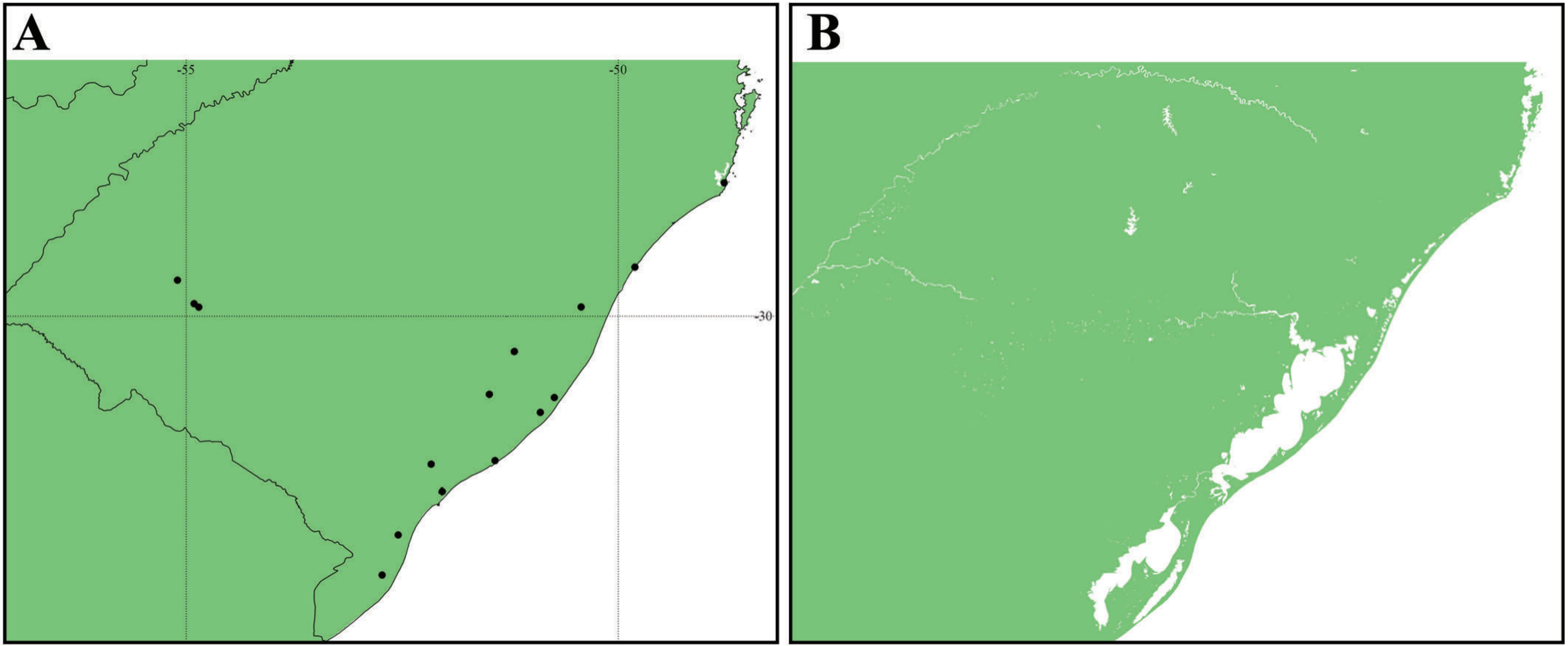
A
B
-55
-50
-30

**Table S1.** Migration estimates obtained with three independent runs of BAYESASS. The values indicate the estimated posterior mean effective migration rate per generation [the fraction of individuals in population *i* (rows) that are migrants derived from population *j* (columns)], and the numbers in parentheses show the standard deviation. Bold values indicate the diagonal (intra-population estimates), and red values indicate the highest migration estimates (those with above zero 95% confidence intervals).

| | I1$_j$ | I2$_j$ | I3$_j$ | W1$_j$ | W2$_j$ | W3$_j$ | N1$_j$ | N2$_j$ | N3$_j$ | S1$_j$ | S2$_j$ | S3$_j$ | S4$_j$ | S5$_j$ | S6$_j$ |
|---|---|---|---|---|---|---|---|---|---|---|---|---|---|---|---|
| I1$_i$ | **0.6850 (0.0180)** | 0.0454 (0.0389) | 0.0179 (0.0171) | 0.0183 (0.0185) | 0.0177 (0.0168) | 0.0330 (0.0280) | 0.0179 (0.0169) | 0.0405 (0.0306) | 0.0175 (0.0164) | 0.0179 (0.0168) | 0.0186 (0.0183) | 0.0181 (0.0172) | 0.0176 (0.0169) | 0.0173 (0.0164) | 0.0174 (0.0164) |
| I2$_i$ | 0.0136 (0.0132) | **0.7980 (0.0339)** | 0.0136 (0.0129) | 0.0134 (0.0131) | 0.0136 (0.0131) | 0.0168 (0.0153) | 0.0136 (0.0130) | 0.0160 (0.0155) | 0.0135 (0.0130) | 0.0173 (0.0159) | 0.0168 (0.0159) | 0.0135 (0.0130) | 0.0135 (0.0131) | 0.0135 (0.0130) | 0.0133 (0.0126) |
| I3$_i$ | 0.0144 (0.0139) | 0.1189 (0.0356) | **0.6806 (0.0134)** | 0.0140 (0.0134) | 0.0211 (0.0185) | 0.0140 (0.0133) | 0.0140 (0.0133) | 0.0232 (0.0199) | 0.0144 (0.0137) | 0.0139 (0.0131) | 0.0146 (0.0142) | 0.0153 (0.0145) | 0.0138 (0.0132) | 0.0140 (0.0132) | 0.0138 (0.0132) |
| W1$_i$ | 0.0175 (0.0169) | 0.0203 (0.0192) | 0.0174 (0.0164) | **0.6842 (0.0165)** | 0.0179 (0.0171) | 0.0189 (0.0178) | 0.0174 (0.0165) | 0.0793 (0.0330) | 0.0177 (0.0167) | 0.0175 (0.0167) | 0.0177 (0.0169) | 0.0174 (0.0168) | 0.0179 (0.0170) | 0.0212 (0.0195) | 0.0176 (0.0167) |
| W2$_i$ | 0.0080 (0.0079) | 0.0101 (0.0094) | 0.0079 (0.0077) | 0.0081 (0.0080) | **0.8792 (0.0257)** | 0.0080 (0.0079) | 0.0080 (0.0079) | 0.0105 (0.0100) | 0.0080 (0.0079) | 0.0087 (0.0085) | 0.0086 (0.0083) | 0.0081 (0.0078) | 0.0081 (0.0078) | 0.0105 (0.0099) | 0.0082 (0.0081) |
| W3$_i$ | 0.0088 (0.0087) | 0.0088 (0.0086) | 0.0088 (0.0087) | 0.0087 (0.0085) | 0.0089 (0.0087) | **0.8694 (0.0270)** | 0.0089 (0.0086) | 0.0098 (0.0095) | 0.0087 (0.0085) | 0.0088 (0.0086) | 0.0087 (0.0087) | 0.0092 (0.0089) | 0.0088 (0.0085) | 0.0149 (0.0124) | 0.0088 (0.0085) |
| N1$_i$ | 0.0185 (0.0175) | 0.0181 (0.0175) | 0.0187 (0.0176) | 0.0186 (0.0173) | 0.0191 (0.0177) | 0.0193 (0.0181) | **0.6849 (0.0171)** | 0.0711 (0.0320) | 0.0187 (0.0177) | 0.0187 (0.0179) | 0.0185 (0.0178) | 0.0203 (0.0194) | 0.0185 (0.0175) | 0.0186 (0.0178) | 0.0183 (0.0175) |
| N2$_i$ | 0.0119 (0.0114) | 0.0119 (0.0113) | 0.0122 (0.0117) | 0.0120 (0.0116) | 0.0138 (0.0134) | 0.0123 (0.0117) | 0.0121 (0.0117) | **0.8201 (0.0329)** | 0.0119 (0.0115) | 0.0129 (0.0126) | 0.0138 (0.0131) | 0.0143 (0.0134) | 0.0119 (0.0116) | 0.0156 (0.0144) | 0.0134 (0.0131) |
| N3$_i$ | 0.0063 (0.0063) | 0.0065 (0.0064) | 0.0064 (0.0064) | 0.0064 (0.0063) | 0.0064 (0.0064) | 0.0065 (0.0063) | 0.0065 (0.0065) | 0.0066 (0.0064) | **0.9055 (0.0216)** | 0.0064 (0.0061) | 0.0086 (0.0086) | 0.0086 (0.0078) | 0.0064 (0.0063) | 0.0064 (0.0063) | 0.0065 (0.0064) |
| S1$_i$ | 0.0132 (0.0127) | 0.0134 (0.0131) | 0.0133 (0.0127) | 0.0136 (0.0134) | 0.0134 (0.0127) | 0.0135 (0.0126) | 0.0133 (0.0129) | 0.0136 (0.0131) | 0.0135 (0.0127) | **0.7483 (0.0703)** | 0.0768 (0.0705) | 0.0136 (0.0130) | 0.0133 (0.0129) | 0.0135 (0.0129) | 0.0138 (0.0133) |
| S2$_i$ | 0.0088 (0.0085) | 0.0087 (0.0084) | 0.0088 (0.0086) | 0.0088 (0.0086) | 0.0088 (0.0085) | 0.0088 (0.0086) | 0.0088 (0.0084) | 0.0088 (0.0085) | 0.0973 (0.0967) | 0.0094 (0.0090) | **0.7873 (0.0966)** | 0.0095 (0.0092) | 0.0088 (0.0086) | 0.0087 (0.0085) | 0.0086 (0.0086) |
| S3$_i$ | 0.0081 (0.0080) | 0.0081 (0.0080) | 0.0081 (0.0079) | 0.0081 (0.0080) | 0.0081 (0.0079) | 0.0087 (0.0083) | 0.0080 (0.0079) | 0.0088 (0.0085) | 0.0098 (0.0094) | 0.0082 (0.0078) | 0.0088 (0.0086) | **0.8814 (0.0253)** | 0.0081 (0.0080) | 0.0085 (0.0083) | 0.0092 (0.0089) |
| S4$_i$ | 0.0122 (0.0117) | 0.0124 (0.0120) | 0.0125 (0.0120) | 0.0125 (0.0123) | 0.0124 (0.0119) | 0.0124 (0.0120) | 0.0122 (0.0118) | 0.0131 (0.0126) | 0.0136 (0.0133) | 0.0125 (0.0119) | 0.0125 (0.0120) | 0.0143 (0.0136) | **0.6792 (0.0121)** | 0.1557 (0.0323) | 0.0125 (0.0120) |
| S5$_i$ | 0.0127 (0.0121) | 0.0125 (0.0122) | 0.0129 (0.0124) | 0.0129 (0.0125) | 0.0127 (0.0121) | 0.0150 (0.0144) | 0.0130 (0.0123) | 0.0131 (0.0126) | 0.0129 (0.0123) | 0.0127 (0.0124) | 0.0129 (0.0124) | 0.0129 (0.0124) | 0.0127 (0.0122) | **0.8180 (0.0322)** | 0.0130 (0.0126) |
| S6$_i$ | 0.0060 (0.0059) | 0.0059 (0.0058) | 0.0059 (0.0058) | 0.0060 (0.0058) | 0.0059 (0.0059) | 0.0061 (0.0060) | 0.0060 (0.0058) | 0.0060 (0.0060) | 0.0059 (0.0058) | 0.0059 (0.0058) | 0.0059 (0.0058) | 0.0061 (0.0060) | 0.0060 (0.0059) | 0.0059 (0.0058) | **0.9164 (0.0193)** |

| | $I1_j$ | $I2_j$ | $I3_j$ | $W1_j$ | $W2_j$ | $W3_j$ | $N1_j$ | $N2_j$ | $N3_j$ | $S1_j$ | $S2_j$ | $S3_j$ | $S4_j$ | $S5_j$ | $S6_j$ |
|---|---|---|---|---|---|---|---|---|---|---|---|---|---|---|---|
| $I1_i$ | **0.6848<br>(0.0172)** | 0.0418<br>(0.0393) | 0.0399<br>(0.0400) | 0.0175<br>(0.0163) | 0.0175<br>(0.0165) | 0.0262<br>(0.0246) | 0.0178<br>(0.0169) | 0.0306<br>(0.0289) | 0.0176<br>(0.0164) | 0.0192<br>(0.0188) | 0.0174<br>(0.0166) | 0.0176<br>(0.0167) | 0.0174<br>(0.0164) | 0.0174<br>(0.0165) | 0.0173<br>(0.0166) |
| $I2_i$ | 0.0135<br>(0.0132) | **0.7976<br>(0.0340)** | 0.0145<br>(0.0139) | 0.0138<br>(0.0135) | 0.0138<br>(0.0131) | 0.0166<br>(0.0153) | 0.0132<br>(0.0126) | 0.0154<br>(0.0146) | 0.0132<br>(0.0126) | 0.0203<br>(0.0179) | 0.0142<br>(0.0134) | 0.0134<br>(0.0129) | 0.0135<br>(0.0131) | 0.0134<br>(0.0127) | 0.0136<br>(0.0129) |
| $I3_i$ | 0.0138<br>(0.0132) | **0.1111<br>(0.0383)** | **0.6858<br>(0.0186)** | 0.0139<br>(0.0133) | 0.0235<br>(0.0197) | 0.0138<br>(0.0131) | 0.0140<br>(0.0138) | 0.0236<br>(0.0202) | 0.0143<br>(0.0137) | 0.0143<br>(0.0137) | 0.0148<br>(0.0140) | 0.0151<br>(0.0144) | 0.0141<br>(0.0136) | 0.0140<br>(0.0134) | 0.0140<br>(0.0134) |
| $W1_i$ | 0.0175<br>(0.0167) | 0.0204<br>(0.0189) | 0.0179<br>(0.0172) | **0.6839<br>(0.0163)** | 0.0173<br>(0.0167) | 0.0192<br>(0.0180) | 0.0176<br>(0.0168) | **0.0805<br>(0.0335)** | 0.0175<br>(0.0170) | 0.0170<br>(0.0163) | 0.0182<br>(0.0175) | 0.0175<br>(0.0166) | 0.0177<br>(0.0169) | 0.0201<br>(0.0185) | 0.0176<br>(0.0167) |
| $W2_i$ | 0.0080<br>(0.0077) | 0.0097<br>(0.0093) | 0.0091<br>(0.0087) | 0.0080<br>(0.0079) | **0.8789<br>(0.0260)** | 0.0082<br>(0.0081) | 0.0080<br>(0.0079) | 0.0101<br>(0.0097) | 0.0081<br>(0.0080) | 0.0093<br>(0.0091) | 0.0082<br>(0.0080) | 0.0081<br>(0.0079) | 0.0080<br>(0.0078) | 0.0104<br>(0.0096) | 0.0079<br>(0.0078) |
| $W3_i$ | 0.0087<br>(0.0086) | 0.0089<br>(0.0086) | 0.0087<br>(0.0085) | 0.0087<br>(0.0084) | 0.0089<br>(0.0086) | **0.8697<br>(0.0270)** | 0.0088<br>(0.0085) | 0.0099<br>(0.0096) | 0.0088<br>(0.0085) | 0.0086<br>(0.0085) | 0.0088<br>(0.0085) | 0.0091<br>(0.0088) | 0.0087<br>(0.0085) | 0.0149<br>(0.0124) | 0.0088<br>(0.0085) |
| $N1_i$ | 0.0186<br>(0.0174) | 0.0186<br>(0.0175) | 0.0187<br>(0.0176) | 0.0181<br>(0.0174) | 0.0186<br>(0.0176) | 0.0189<br>(0.0181) | **0.6854<br>(0.0178)** | **0.0713<br>(0.0319)** | 0.0182<br>(0.0171) | 0.0188<br>(0.0176) | 0.0184<br>(0.0175) | 0.0202<br>(0.0190) | 0.0185<br>(0.0177) | 0.0188<br>(0.0180) | 0.0188<br>(0.0178) |
| $N2_i$ | 0.0118<br>(0.0114) | 0.0120<br>(0.0118) | 0.0119<br>(0.0116) | 0.0118<br>(0.0113) | 0.0138<br>(0.0132) | 0.0124<br>(0.0119) | 0.0116<br>(0.0114) | **0.8204<br>(0.0323)** | 0.0121<br>(0.0115) | 0.0135<br>(0.0128) | 0.0137<br>(0.0131) | 0.0143<br>(0.0137) | 0.0119<br>(0.0115) | 0.0154<br>(0.0144) | 0.0134<br>(0.0128) |
| $N3_i$ | 0.0065<br>(0.0064) | 0.0064<br>(0.0062) | 0.0065<br>(0.0065) | 0.0064<br>(0.0063) | 0.0066<br>(0.0063) | 0.0064<br>(0.0063) | 0.0064<br>(0.0063) | 0.0066<br>(0.0064) | **0.9034<br>(0.0216)** | 0.0064<br>(0.0064) | 0.0102<br>(0.0097) | 0.0090<br>(0.0080) | 0.0065<br>(0.0064) | 0.0064<br>(0.0063) | 0.0064<br>(0.0062) |
| $S1_i$ | 0.0134<br>(0.0128) | 0.0133<br>(0.0126) | 0.0132<br>(0.0126) | 0.0132<br>(0.0127) | 0.0133<br>(0.0127) | 0.0135<br>(0.0132) | 0.0135<br>(0.0130) | 0.0138<br>(0.0130) | 0.0132<br>(0.0127) | **0.8033<br>(0.0452)** | 0.0219<br>(0.0358) | 0.0135<br>(0.0130) | 0.0135<br>(0.0129) | 0.0133<br>(0.0129) | 0.0140<br>(0.0133) |
| $S2_i$ | 0.0087<br>(0.0085) | 0.0090<br>(0.0087) | 0.0088<br>(0.0086) | 0.0088<br>(0.0087) | 0.0088<br>(0.0084) | 0.0089<br>(0.0087) | 0.0086<br>(0.0084) | 0.0088<br>(0.0086) | 0.0244<br>(0.0399) | 0.0097<br>(0.0093) | **0.8593<br>(0.0457)** | 0.0096<br>(0.0093) | 0.0088<br>(0.0085) | 0.0088<br>(0.0087) | 0.0090<br>(0.0087) |
| $S3_i$ | 0.0082<br>(0.0079) | 0.0081<br>(0.0079) | 0.0082<br>(0.0080) | 0.0083<br>(0.0080) | 0.0081<br>(0.0079) | 0.0088<br>(0.0084) | 0.0082<br>(0.0079) | 0.0089<br>(0.0086) | 0.0097<br>(0.0092) | 0.0082<br>(0.0081) | 0.0096<br>(0.0092) | **0.8802<br>(0.0255)** | 0.0081<br>(0.0079) | 0.0086<br>(0.0083) | 0.0090<br>(0.0086) |
| $S4_i$ | 0.0123<br>(0.0119) | 0.0123<br>(0.0119) | 0.0122<br>(0.0118) | 0.0123<br>(0.0117) | 0.0123<br>(0.0119) | 0.0122<br>(0.0116) | 0.0123<br>(0.0116) | 0.0133<br>(0.0126) | 0.0135<br>(0.0131) | 0.0123<br>(0.0118) | 0.0129<br>(0.0125) | 0.0144<br>(0.0137) | **0.6790<br>(0.0117)** | **0.1564<br>(0.0318)** | 0.0124<br>(0.0120) |
| $S5_i$ | 0.0130<br>(0.0125) | 0.0128<br>(0.0125) | 0.0128<br>(0.0121) | 0.0128<br>(0.0124) | 0.0130<br>(0.0125) | 0.0149<br>(0.0141) | 0.0126<br>(0.0122) | 0.0129<br>(0.0123) | 0.0128<br>(0.0122) | 0.0129<br>(0.0125) | 0.0132<br>(0.0128) | 0.0128<br>(0.0124) | 0.0126<br>(0.0120) | **0.8182<br>(0.0322)** | 0.0128<br>(0.0123) |
| $S6_i$ | 0.0059<br>(0.0059) | 0.0060<br>(0.0058) | 0.0060<br>(0.0059) | 0.0059<br>(0.0058) | 0.0059<br>(0.0058) | 0.0061<br>(0.0060) | 0.0060<br>(0.0060) | 0.0061<br>(0.0060) | 0.0061<br>(0.0060) | 0.0060<br>(0.0058) | 0.0060<br>(0.0058) | 0.0062<br>(0.0062) | 0.0060<br>(0.0057) | 0.0061<br>(0.0061) | **0.9157<br>(0.0194)** |

| | $I1_j$ | $I2_j$ | $I3_j$ | $W1_j$ | $W2_j$ | $W3_j$ | $N1_j$ | $N2_j$ | $N3_j$ | $S1_j$ | $S2_j$ | $S3_j$ | $S4_j$ | $S5_j$ | $S6_j$ |
|---|---|---|---|---|---|---|---|---|---|---|---|---|---|---|---|
| $I1_i$ | **0.6852**<br>**(0.0185)** | 0.0557<br>(0.0411) | 0.0219<br>(0.0246) | 0.0177<br>(0.0170) | 0.0175<br>(0.0167) | 0.0259<br>(0.0244) | 0.0178<br>(0.0169) | 0.0338<br>(0.0299) | 0.0174<br>(0.0164) | 0.0187<br>(0.0180) | 0.0180<br>(0.0177) | 0.0175<br>(0.0167) | 0.0177<br>(0.0167) | 0.0178<br>(0.0171) | 0.0175<br>(0.0164) |
| $I2_i$ | 0.0134<br>(0.0130) | **0.7989**<br>**(0.0338)** | 0.0139<br>(0.0132) | 0.0135<br>(0.0128) | 0.0136<br>(0.0131) | 0.0171<br>(0.0158) | 0.0134<br>(0.0129) | 0.0156<br>(0.0150) | 0.0133<br>(0.0130) | 0.0179<br>(0.0165) | 0.0156<br>(0.0149) | 0.0133<br>(0.0127) | 0.0135<br>(0.0128) | 0.0135<br>(0.0130) | 0.0135<br>(0.0131) |
| $I3_i$ | 0.0143<br>(0.0136) | **0.1194**<br>**(0.0364)** | **0.6813**<br>**(0.0143)** | 0.0139<br>(0.0133) | 0.0211<br>(0.0188) | 0.0139<br>(0.0132) | 0.0137<br>(0.0133) | 0.0222<br>(0.0196) | 0.0143<br>(0.0138) | 0.0142<br>(0.0137) | 0.0146<br>(0.0141) | 0.0150<br>(0.0144) | 0.0139<br>(0.0133) | 0.0141<br>(0.0135) | 0.0140<br>(0.0135) |
| $W1_i$ | 0.0176<br>(0.0167) | 0.0207<br>(0.0194) | 0.0174<br>(0.0165) | **0.6845**<br>**(0.0168)** | 0.0176<br>(0.0170) | 0.0185<br>(0.0175) | 0.0175<br>(0.0168) | **0.0801**<br>**(0.0331)** | 0.0177<br>(0.0167) | 0.0176<br>(0.0168) | 0.0177<br>(0.0169) | 0.0175<br>(0.0165) | 0.0174<br>(0.0162) | 0.0209<br>(0.0192) | 0.0172<br>(0.0164) |
| $W2_i$ | 0.0081<br>(0.0079) | 0.0099<br>(0.0093) | 0.0081<br>(0.0079) | 0.0082<br>(0.0078) | **0.8791**<br>**(0.0253)** | 0.0081<br>(0.0079) | 0.0080<br>(0.0079) | 0.0104<br>(0.0099) | 0.0080<br>(0.0079) | 0.0088<br>(0.0085) | 0.0083<br>(0.0081) | 0.0083<br>(0.0081) | 0.0080<br>(0.0078) | 0.0104<br>(0.0097) | 0.0082<br>(0.0079) |
| $W3_i$ | 0.0088<br>(0.0086) | 0.0088<br>(0.0085) | 0.0087<br>(0.0086) | 0.0088<br>(0.0087) | 0.0087<br>(0.0086) | **0.8698**<br>**(0.0270)** | 0.0089<br>(0.0084) | 0.0097<br>(0.0092) | 0.0088<br>(0.0086) | 0.0087<br>(0.0084) | 0.0088<br>(0.0086) | 0.0091<br>(0.0086) | 0.0087<br>(0.0083) | 0.0148<br>(0.0122) | 0.0088<br>(0.0086) |
| $N1_i$ | 0.0185<br>(0.0178) | 0.0186<br>(0.0178) | 0.0183<br>(0.0172) | 0.0184<br>(0.0175) | 0.0188<br>(0.0178) | 0.0194<br>(0.0185) | **0.6848**<br>**(0.0173)** | **0.0720**<br>**(0.0326)** | 0.0183<br>(0.0172) | 0.0183<br>(0.0172) | 0.0188<br>(0.0179) | 0.0202<br>(0.0187) | 0.0185<br>(0.0177) | 0.0187<br>(0.0177) | 0.0183<br>(0.0175) |
| $N2_i$ | 0.0120<br>(0.0115) | 0.0120<br>(0.0115) | 0.0117<br>(0.0114) | 0.0119<br>(0.0117) | 0.0138<br>(0.0134) | 0.0123<br>(0.0118) | 0.0119<br>(0.0116) | **0.8204**<br>**(0.0323)** | 0.0121<br>(0.0117) | 0.0132<br>(0.0128) | 0.0139<br>(0.0133) | 0.0141<br>(0.0132) | 0.0119<br>(0.0116) | 0.0154<br>(0.0145) | 0.0133<br>(0.0128) |
| $N3_i$ | 0.0065<br>(0.0065) | 0.0064<br>(0.0063) | 0.0064<br>(0.0063) | 0.0065<br>(0.0064) | 0.0065<br>(0.0064) | 0.0064<br>(0.0062) | 0.0065<br>(0.0063) | 0.0065<br>(0.0065) | **0.9046**<br>**(0.0216)** | 0.0065<br>(0.0063) | 0.0091<br>(0.0089) | 0.0086<br>(0.0079) | 0.0064<br>(0.0063) | 0.0065<br>(0.0063) | 0.0066<br>(0.0064) |
| $S1_i$ | 0.0132<br>(0.0126) | 0.0133<br>(0.0127) | 0.0133<br>(0.0127) | 0.0132<br>(0.0127) | 0.0136<br>(0.0129) | 0.0134<br>(0.0129) | 0.0134<br>(0.0126) | 0.0135<br>(0.0132) | 0.0133<br>(0.0127) | **0.7636**<br>**(0.0693)** | 0.0623<br>(0.0674) | 0.0136<br>(0.0130) | 0.0132<br>(0.0126) | 0.0133<br>(0.0130) | 0.0139<br>(0.0133) |
| $S2_i$ | 0.0087<br>(0.0085) | 0.0088<br>(0.0086) | 0.0088<br>(0.0087) | 0.0087<br>(0.0085) | 0.0086<br>(0.0084) | 0.0090<br>(0.0087) | 0.0087<br>(0.0084) | 0.0089<br>(0.0087) | 0.0836<br>(0.0938) | 0.0092<br>(0.0089) | **0.8010**<br>**(0.0933)** | 0.0092<br>(0.0089) | 0.0087<br>(0.0084) | 0.0089<br>(0.0090) | 0.0090<br>(0.0087) |
| $S3_i$ | 0.0081<br>(0.0078) | 0.0083<br>(0.0081) | 0.0081<br>(0.0079) | 0.0083<br>(0.0080) | 0.0082<br>(0.0079) | 0.0088<br>(0.0086) | 0.0083<br>(0.0081) | 0.0088<br>(0.0085) | 0.0098<br>(0.0095) | 0.0080<br>(0.0080) | 0.0090<br>(0.0089) | **0.8810**<br>**(0.0253)** | 0.0081<br>(0.0079) | 0.0084<br>(0.0083) | 0.0089<br>(0.0086) |
| $S4_i$ | 0.0123<br>(0.0116) | 0.0125<br>(0.0120) | 0.0123<br>(0.0118) | 0.0125<br>(0.0120) | 0.0124<br>(0.0118) | 0.0124<br>(0.0117) | 0.0121<br>(0.0117) | 0.0128<br>(0.0123) | 0.0132<br>(0.0125) | 0.0125<br>(0.0121) | 0.0129<br>(0.0125) | 0.0145<br>(0.0136) | **0.6792**<br>**(0.0121)** | **0.1560**<br>**(0.0322)** | 0.0124<br>(0.0122) |
| $S5_i$ | 0.0129<br>(0.0126) | 0.0127<br>(0.0123) | 0.0128<br>(0.0123) | 0.0129<br>(0.0124) | 0.0129<br>(0.0124) | 0.0150<br>(0.0142) | 0.0129<br>(0.0124) | 0.0130<br>(0.0127) | 0.0129<br>(0.0123) | 0.0127<br>(0.0122) | 0.0130<br>(0.0124) | 0.0130<br>(0.0126) | 0.0128<br>(0.0121) | **0.8176**<br>**(0.0324)** | 0.0129<br>(0.0124) |
| $S6_i$ | 0.0059<br>(0.0058) | 0.0060<br>(0.0059) | 0.0059<br>(0.0058) | 0.0061<br>(0.0060) | 0.0059<br>(0.0060) | 0.0060<br>(0.0058) | 0.0060<br>(0.0059) | 0.0060<br>(0.0060) | 0.0059<br>(0.0058) | 0.0058<br>(0.0058) | 0.0061<br>(0.0059) | 0.0060<br>(0.0059) | 0.0059<br>(0.0059) | 0.0059<br>(0.0057) | **0.9165**<br>**(0.0192)** |
